# Temporal Biodynamics: An AI Platform for Identification of Stage-Relevant Targets and Biomarkers

**DOI:** 10.64898/2026.06.03.729984

**Authors:** Parth Natekar, Baixue Yao, Sara Mohammad-Taheri, Anna Rusnak, Nicolas A. Gort-Freitas, Jonathan Fillatre, Joseph J Raymond, Sachit D. Saksena, Scott Lipnick, Artem Sokolov

## Abstract

Temporal modeling of disease progression is poised to revolutionize the process of target identification, leading to better characterization of and intervention at the critical early stages of chronic conditions. Temporal Biodynamics is an artificial intelligence-driven platform that leverages within-tissue heterogeneity in cross-sectional cohorts to assemble a single, continuous trajectory of transcriptomic changes between health and disease. We demonstrate that the platform enriches for known disease-associated genes and proteins by more than 50% over the conventional case-control comparisons. When compared to other published pseudotime methods, our models were better at extracting disease-relevant signals in the presence of confounders and co-morbidities. The Temporal Biodynamics platform enables rich profiling of a disease continuum, providing temporal insights that are otherwise hidden by the traditional discrete staging of chronic diseases. This includes detecting cascades of molecular events, providing clues regarding causality, and increasing confidence in blood-based protein biomarkers using tissue-based context.

## Introduction

Inadequate efficacy drives over 50% of late-stage drug development failures, in many cases due to the wrong target, patient population, or stage of intervention^1,2^. Novel target identification is typically focused on comparing molecular measurements (e.g., RNA sequencing) of diseased cohorts to those of healthy counterparts. Such case-control comparisons fail to capture the nuances of disease progression, which is inherently continuous and is driven by temporal transitions in the underlying biological processes. This is particularly prominent in chronic diseases, where discrete diagnostic staging is traditionally used to categorize patients into distinct clinical groups. While categorical frameworks have proven useful for clinical decision-making, they fundamentally misrepresent the underlying biology of disease progression. Recent advances in single-cell genomics have repeatedly revealed that chronic diseases involve a continuum of cellular states, with individual cells transitioning gradually from healthy to diseased phenotypes rather than occupying a small number of discrete molecular states^3–7^.

Single-cell RNA sequencing (scRNA-seq) has emerged as a powerful technology for characterizing cellular heterogeneity of diseased tissue, enabling the profiling of thousands to millions of individual cells and revealing that each tissue specimen represents a heterogeneous mixture of cells at different points along the disease continuum. To better understand continuous cellular transitions, single-cell data has been the focus of numerous pseudotime and trajectory inference methods^5,8–11^ . These methods typically operate along the main axes of variation and are primarily applied to developmental biology, where differentiation processes dominate the transcriptional signal, and longitudinal sampling or time-series data can capture cells across the entire developmental trajectory. In contrast, most large-scale disease studies contain a single sample per patient, providing a static snapshot of the cellular composition in a population. Unlike model systems where a single organism progresses through all phases of development, disease progression must be reconstructed from overlapping and incomplete cellular distributions sampled from patients at different clinical stages. Critically, disease labels are available only at the patient level rather than for individual cells, making them inextricably linked to other patient-level covariates (e.g., age, sex, medication usage, environmental exposure, comorbidities) and technical batch effects. This introduces the risk of confounding and makes it difficult to distinguish true disease-associated cellular variation from unrelated factors. The cross-sectional data structure, combined with patient-level labeling and the fact that disease-associated transcriptional shifts are considerably more subtle than developmental biology, creates a challenging inference problem for traditional pseudotime methods.

In this work, we present Temporal Biodynamics, an artificial intelligence-driven platform to assemble a continuous trajectory of transcriptomic changes between health and disease using cross-sectional tissue specimens. The platform makes use of cell state heterogeneity found within diseased tissues of individual patients, which enables stitching of overlapping cell states from neighboring clinical stages. Unlike existing pseudotime trajectory modeling, Temporal Biodynamics uses clinical staging information to orient the data and extract subtle disease signals that may otherwise be masked by confounding variables such as comorbidities. The time-based nature of the platform enables rich profiling of a disease continuum that is not possible with conventional case-control comparisons. This includes detecting cascades of molecular events, providing clues regarding causality, and characterizing disease in its earliest stages, before symptoms appear. Taken together, these capabilities enable the identification of novel, stage-relevant targets and biomarkers that are otherwise hidden from conventional approaches.

## Results

### Overview of the Temporal Biodynamics platform

The Temporal Biodynamics platform requires several assumptions to support effective application to a disease area. First, disease progression must represent a continuum of cellular states rather than a discrete set (e.g., as conferred by the accumulation of mutations in cancer). Second, each patient sample must contain a heterogeneous mixture of cells at different disease states, and cells from patients at a more advanced clinical stage are expected, on average, to be further along the disease continuum than cells from healthier patients. Third, cells from different patients must share a common underlying molecular disease trajectory, such that cells at equivalent disease states exhibit transcriptional similarity that is independent of patient identity. The shared trajectory may be obscured by patient-specific confounders, including biological factors (e.g., age, sex, genetics, comorbidities) and technical factors (batch effects). Given these assumptions, our platform generates Temporal Biodynamic Representations (TBRs) of disease by aligning overlapping cellular distributions across patients and using patient-level clinical label supervision to arrive at a coherent trajectory spanning the full disease spectrum (Figure 1A).

**Figure 1:**
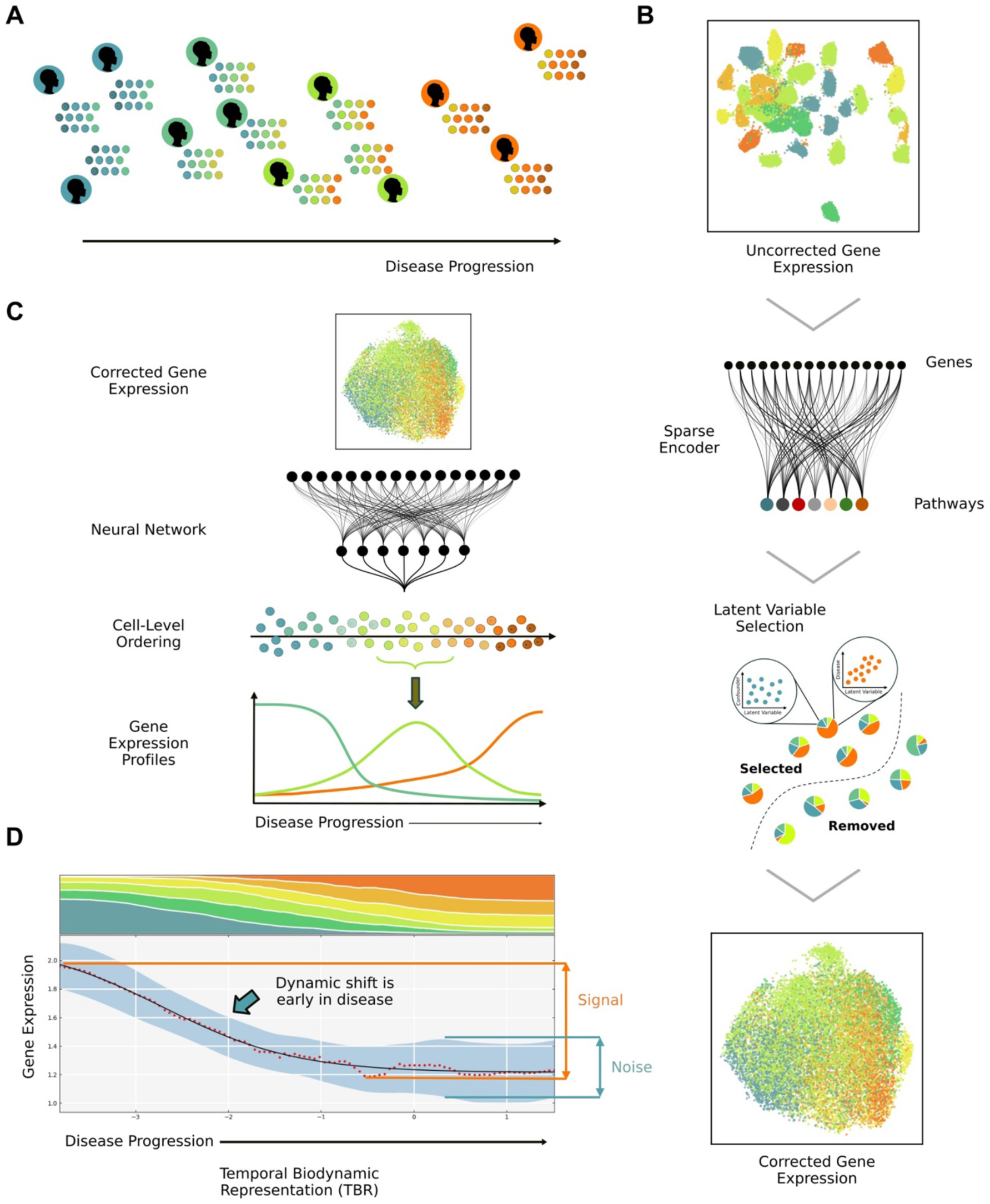
Method overview. **(A)** A schematic representation of single-cell sequencing data for sample tissue from a cohort of patients ordered from lowest to highest disease severity, using patient-level clinical labels as guidance. Colors encode disease severity at the level of both cells and patients, with blue representing healthy or least diseased, green/yellow corresponding to early/mid disease stages, and orange representing late stages. The diversity of colors for cells associated with each patient represents within-tissue heterogeneity of cell states. **(B)** Uniform Manifold Approximation and Projection (UMAP) plots showing uncorrected gene expression data for a real-world patient cohort (top-most), and latent autoencoder (AE) space after the confounding axes of variation were removed (bottom-most). Points represent individual cells, colored by disease severity. The middle panel shows the architecture of the autoencoder used to project the original gene expression data into a lower-dimensional representation. Pathway databases are used to define connections between layers. Axes of variation in the lower-dimensional representation are excluded based on their correlation with disease labels vs. their ability to distinguish individual patients. **(C)** An ordinal neural network model is trained on the corrected data to order cells along a single continuum of disease using patient-level supervision. Profiles of expression for each gene are computed by convolving a kernel against gene expression of individual ordered cells. An example Temporal Biodynamic Representation (TBR) is shown as the output. The x axis represents the disease continuum with the streamgraph at the top showing the proportion of cells from each disease condition that were mapped to the corresponding region along the continuum. The plot below shows the change in expression (y axis) of a single gene along the disease continuum. The red dots represent values from the kernel convolution. The black line is a double-sigmoid fit to those points, and the shaded blue region corresponds to the standard deviation.

TBR construction is performed for each cell type separately, enabling a deeper understanding of cell state transitions from a cross-sectional sampling of disease conditions. A multi-stage workflow separates disease-relevant transitions from potential confounder effects (Figure 1B,C). Following cell type annotation and quality control assessment, high-dimensional single-cell gene expression data is first projected into a lower-dimensional latent space using PathwayAE, a constrained autoencoder (AE) that maps each latent dimension to a known biological pathway (see Methods). Critically, PathwayAE identifies and discards latent dimensions that align more strongly with known confounders than with disease labels, amplifying disease-specific signals that would otherwise be obscured by patient-specific effects (Figure 1B). We find that this step is especially useful in low (<100) patient number scenarios, where the data is insufficient to properly capture patient-to-patient variability among the combinatoric complexity of real-world confounders. In addition to removing confounder associations, the pathway-constrained architecture of PathwayAE ensures that each remaining latent dimension is interpretable in the context of the corresponding biological pathway.

Given patient-informed embeddings from PathwayAE, a fully-connected ordinal-regression neural network is trained to arrange individual cells along a shared one-dimensional disease axis, thereby reconstructing the full disease continuum from overlapping but incomplete cellular distributions given by patients at different clinical stages (Figure 1C). The ordinal regression aspect of the model aims to place cells from patients at more severe clinical stages further along the disease continuum, while the overall learning task ensures that transcriptionally similar cells end up with the same coordinate along the disease severity axis.

After arranging cells along the disease continuum, gene expression profiles are constructed by applying kernel smoothing to single-cell measurements, with bootstrap resampling providing confidence intervals. Finally, a signal-to-noise ratio is computed from the overall gene dynamics across the disease continuum with noise capturing local expression variability (Figure 1C). Statistical significance is assessed by comparing the observed gene dynamics to null distributions generated from shuffling patient labels across multiple parallel models, explicitly testing whether variation disappears when meaningful clinical labels are removed.

### TBRs recover true cell-level trajectories from simulated disease data

To rigorously evaluate TBR performance under controlled conditions, we constructed a simulated disease dataset from a time series characterizing the reprogramming of mouse embryonic fibroblasts to induced endoderm progenitor (iEP) cells^12^ . Cells were ordered by their continuous labels representing the true state of differentiation. Each simulated patient was constructed by sampling the differentiation continuum with a Gaussian distribution centered at a particular cell state. Cell-level labels in the resulting mixtures were discretized into distinct “disease stages” and converted to patient-level “clinical” labels by a majority vote (Figure 2A). To recapitulate real-world challenges, we added sample-level technical noise and introduced confounding effects in the form of simulated batch effects and biological covariates. The resulting simulated dataset faithfully reflects the fundamental constraints of chronic disease modeling from cross-sectional snapshots: (1) each sample contains a distribution of cells at varying disease states, (2) the distribution of cell states is centered around, but not restricted to, the patient-level clinical label, and (3) disease signals are obscured by patient-specific confounders and technical noise.

**Figure 2:**
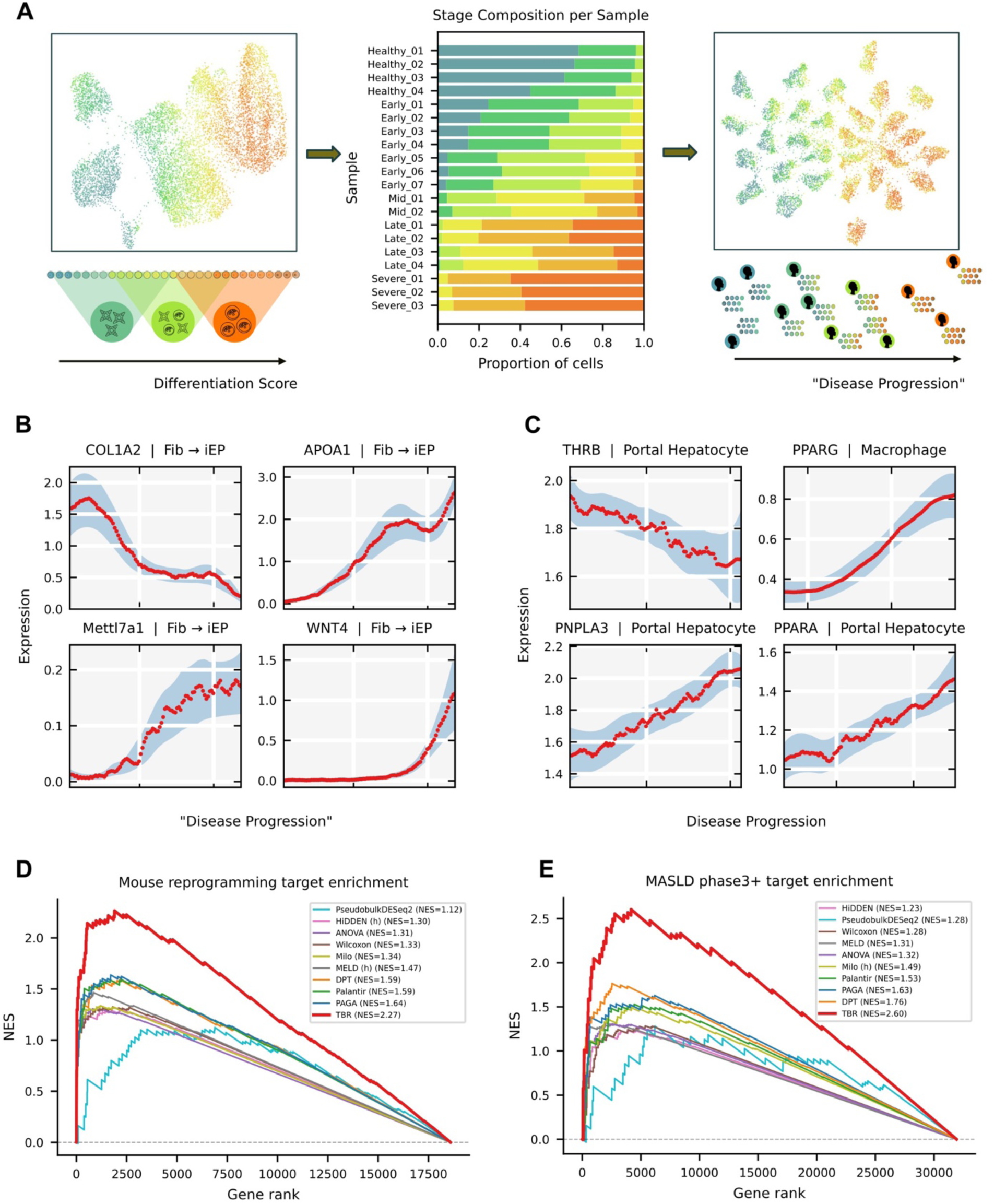
Comparison of TBRs against traditional differential expression methods and pseudo-time trajectory modeling. **(A)** Left: A published dataset characterizing the reprogramming of mouse embryonic fibroblast (MEF) to induced endoderm progenitors (iEP) captures the true level of differentiation using continuous labels for each cell. Middle: A set of simulated patient samples is created using mixtures of cells at various stages along the differentiation trajectory. Right: Patient-level confounding noise is added to the mixtures, creating a dataset that mimics real-world snapshot data of patients at different chronic disease stages with within-tissue molecular heterogeneity and patient-to-patient variability. **(B)** TBR-derived profiles showing gene expression changes along “disease progression” (reprogramming axis) for the mouse fibroblast-to-iEP dataset. Red dots correspond to average expression levels computed by convolving single-cell measurements with a kernel. The shaded blue regions are standard deviations. **(C)** TBR expression profiles for genes in Phase III+ clinical trials for MASLD. The plot elements are as in panel (B). The cell type used for constructing each TBR model is included in the corresponding plot title. **(D, E)** Standard Gene Set Enrichment Analysis (GSEA) curves showing the ability of each method to rank genes known to be associated with the reprogramming of MEF towards iEP (D) and targets of drugs that have reached Phase III+ clinical trials for MASLD (E).

We first assessed whether TBRs could reconstruct true continuous cell differentiation scores from the discrete “clinical” labels given to heterogeneous mixtures in the simulated patient population. We also asked whether the reconstructed trajectories recapitulate known cell differentiation biology. During the initial evaluation, we found that the gene expression profiles constructed by TBR models show reprogramming markers changing in a temporally consistent manner (Figure 2B). Specifically, a canonical fibroblast marker *Col1a2* is downregulated immediately, marking rapid exit from somatic identity. A hepatic identity marker *Apoa1* is upregulated from the earliest stages and sustained throughout the trajectory. Mid-trajectory upregulation was observed for *Mettl7a1*, consistent with its role at the Day 21 reprogramming bifurcation. Lastly, *Wnt4* upregulation is restricted to the late trajectory, coinciding with the final transition to the fully reprogrammed iEP state.

We next compared TBR performance against unsupervised trajectory inference methods (PAGA, DPT, Palantir)^8,9,13^ and supervised label refinement approaches (HiDDEN, MELD, Milo)^11,14,15^ . Standard approaches were also included in the comparison, both as binary case-control contrasts (Wilcoxon) and as multi-stage ANOVA. TBRs achieved the highest correlation with true labels (r = 0.95, p < 0.001) and concordance index (CI = 0.89) across all methods tested (Table 1; Suppl. Figure 1). Unsupervised trajectory methods showed substantially lower performance (r = [0.35, 0.55], CI = [0.62, 0.75]), confirming that approaches operating along dominant axes of variation are misled by noise and confounders and fail to recover the true underlying disease trajectory. Supervised and label refinement methods showed better performance at capturing the underlying disease axis (r = [0.81, 0.82], CI = [0.70, 0.82]) but were still inadequate for reconstructing the original continuum of molecular states.

**Table 1:** Performance of TBR models compared to other trajectory and categorical contrast methods using a dataset characterizing the reprogramming of mouse fibroblasts to induced endoderm progenitors (iEP). The highest performing implementation for each method is shown, with the corresponding dimensionality reduction/batch correction method mentioned in brackets. Known Gene Enrichment reports normalized enrichment scores from the standard gene set enrichment analysis evaluating the ability of each method to find the curated list of genes known to be associated with the cell differentiation processes. Disease State Concordance and Correlation report the ability of each method to reconstruct the original order of differentiation cell states. Transcriptional Continuity reports the average transcriptional similarity of neighboring cells along the reconstructed disease continuum.

| Method FDSF | Known Gene Enrichment | Disease State Concordance | Disease State Correlation | Transcriptional Continuity |
| --- | --- | --- | --- | --- |
| **TBR** | **2.26** | **0.89** | **0.95** | **0.98** |
| ANOVA | 1.31 | NA | NA | NA |
| Wilcoxon | 1.21 | NA | NA | NA |
| Pseudobulk DESeq2 | 1.12 | NA | NA | NA |
| PAGA (PCA) | 1.63 | 0.64 | 0.35 | 0.58 |
| Palantir (PCA) | 1.59 | 0.75 | 0.55 | -0.14 |
| DPT (PCA) | 1.59 | 0.62 | 0.35 | 0.15 |
| Milo (PCA) | 1.33 | 0.70 | 0.81 | 0.63 |
| MELD (Harmony) | 1.46 | 0.82 | 0.82 | 0.90 |
| HIDDEN (Harmony) | 1.30 | 0.81 | 0.82 | 0.74 |

### TBRs outperform existing methods at identifying disease-associated genes

We next evaluated whether accurate trajectory reconstruction translates to improved identification of disease-relevant genes. Using known reprogramming factors and key regulators from the original study^12^ as true disease-associated genes, we computed gene set enrichment scores^16^ for gene rankings produced by each method. TBRs achieved significantly higher enrichment scores (NES = 2.265, p < 0.001) compared to all approaches (Figure 2D, Table 1). Traditional methods that rely on bulk differential expression (PseudobulkDESeq2) or categorical comparisons (Wilcoxon, ANOVA) achieved the lowest enrichment, highlighting the challenges of identifying relevant biological processes without leveraging within-sample heterogeneity. Unsupervised methods showed intermediate enrichment, reflecting their inability to distinguish disease-relevant variation from technical noise and confounders. Label refinement methods also showed low to intermediate enrichment, confirming that while they identify disease-affected cells, the subsequent collapse to discrete groups for differential expression gene ranking loses critical continuous gradient information. Lastly, we measured transcriptional continuity of single cell trajectories to assess alignment with the expected underlying biology of smooth cell state transitions. TBR and MELD were observed to yield the smoothest models.

Having validated TBR performance on a simulated cohort constructed from cell differentiation data with known ground truth, we extended the analysis to a real-world dataset covering metabolically associated steatotic liver disease (MASLD) and metabolic dysfunction-associated steatohepatitis (MASH) patients. The dataset was constructed by performing single-nuclei sequencing on 44 liver specimens obtained from patients who underwent bariatric surgery (see Methods). The specimens were reviewed by a certified pathologist to establish patient-level clinical stages spanning healthy individuals and steatotic disease both in the presence and absence of fibrosis. To establish ground truth in the absence of cell-level labels, we employed two complementary validation strategies. First, we manually curated gene lists from clinical literature and genetic studies to capture key known disease processes (e.g., steatosis, inflammation, fibrosis, metabolic dysfunction). Second, we identified genes targeted by drugs that entered phase III clinical trials for MASLD/MASH.

The overall performance trends were similar to those in the simulated murine embryonic fibroblast dataset, with TBRs showing significantly higher enrichment for both gene sets compared to all other methods (Figure 2E, Table 2). The dual validation approach demonstrates that TBRs recover genes that hold both biological relevance (literature-curated genes) and translational potential (clinical trial targets). In particular, we observed that key disease markers targeted in clinical trials show meaningful changes in TBR-derived gene expression profiles (Figure 2C): *THRB* (target of the agonist resmetirom) is consistently downregulated in hepatocytes, reflecting loss of metabolic control; *PPARA* and *PPARG* (targets of agonist lanifibranor) are upregulated, consistent with a compensatory inflammatory response^17–19^; and *PNPLA3* (target of RNAi ALN-PNP) is consistently upregulated, reflecting increased pathogenic lipid remodeling.

**Table 2:**
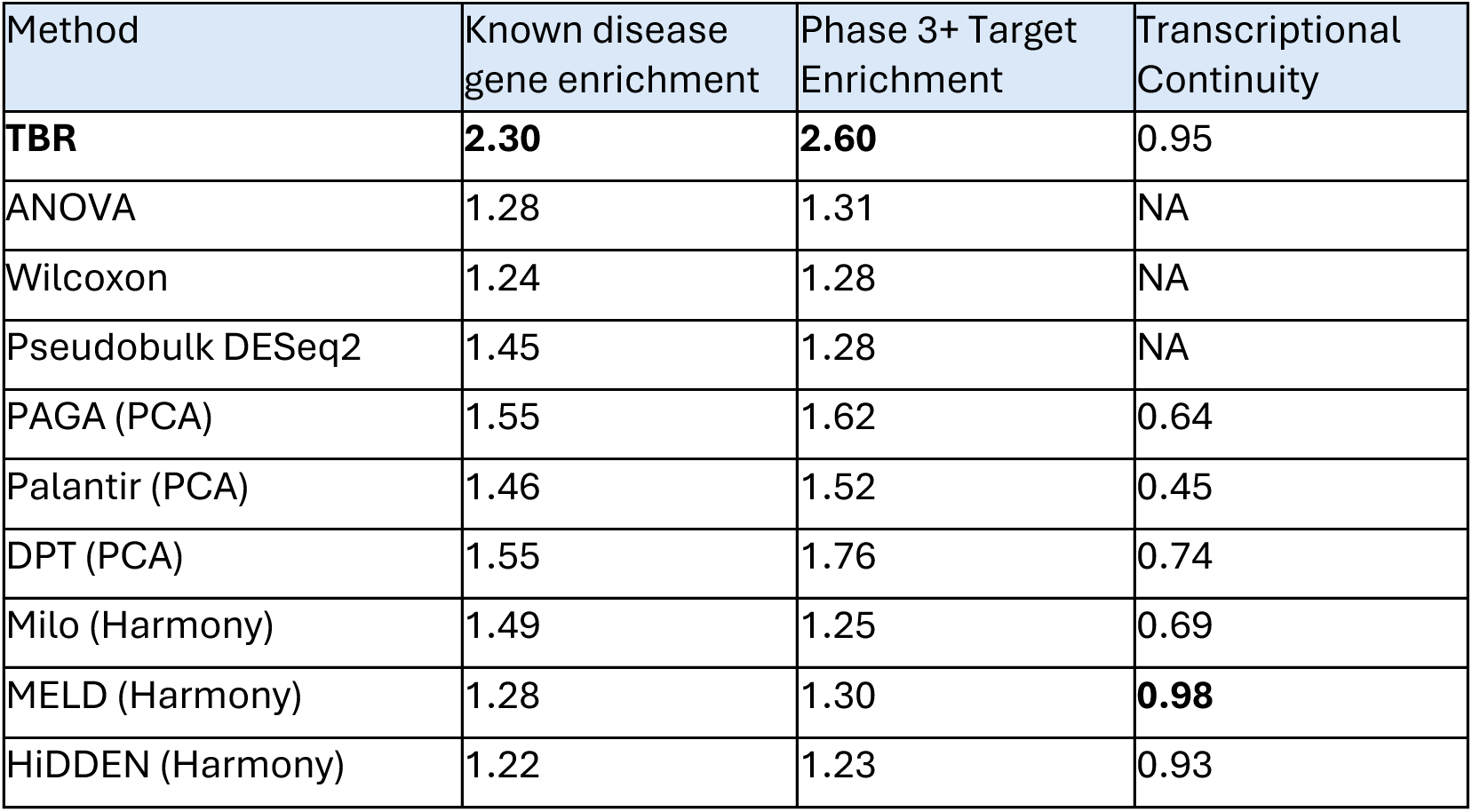
Performance of TBR models compared to other trajectory and categorical contrast methods on a real-world dataset capturing MASLD/MASH disease progression. All elements are as in Table 1, with the only distinction that gene set enrichment scores are reported separately for the literature-curated set and the targets of drugs that entered Phase III clinical trials.

### TBRs identify genes with high within-patient disease trajectory concordance

Target discovery often suffers from high attrition and false-positive rates because standard comparisons of gene expression levels across patients capture demographic noise and systemic variations more readily than true disease biology^20^ . However, genes whose expression changes correlate with disease pathology *within* a single patient have the perfect “internal control”, because the patient’s genetic and systemic background is constant for all their cells. If a gene’s expression correlates to localized disease pathology across multiple independent patients, it increases the probability that the gene has a true association with disease progression and not an unrelated (possibly unmeasured) confounding factor. By contrasting gene expression changes of individual patients against the overall expression profile, the Temporal Biodynamic platform can lower the false discovery rate and nominate therapeutic targets that are better aligned with disease biology.

To quantify intra-patient concordance, we define the Directionally Similar Fraction (DSF) to be the fraction of patients for whom a gene changes in the same direction within their cells along the disease trajectory as the overall population-level trend. The metric introduces a novel dimension for distinguishing gene expression trends (Figure 3A). For example, both Insulin-like Growth Factor 1 (*IGF1*) and Pre-B-cell leukemia transcription factor 1 (*PBX1*) appear to correlate with disease severity in standard bulk or pseudo-bulk analyses. However, the high DSF value for *IGF1* reveals that the gene has a continuous decrease in expression as individual cells transition from healthy to MASH F2, and that the trend is consistent within most patients. Conversely, the low DSF value of *PBX1* captures the fact that its expression correlates with disease across the entire population but lacks a consistent, continuous gradient at the level of individual patients. This suggests that *PBX1* variance may be driven by patient-to-patient variability rather than a genuine association with disease state, while *IGF1* closely aligns with disease biology.

**Figure 3:**
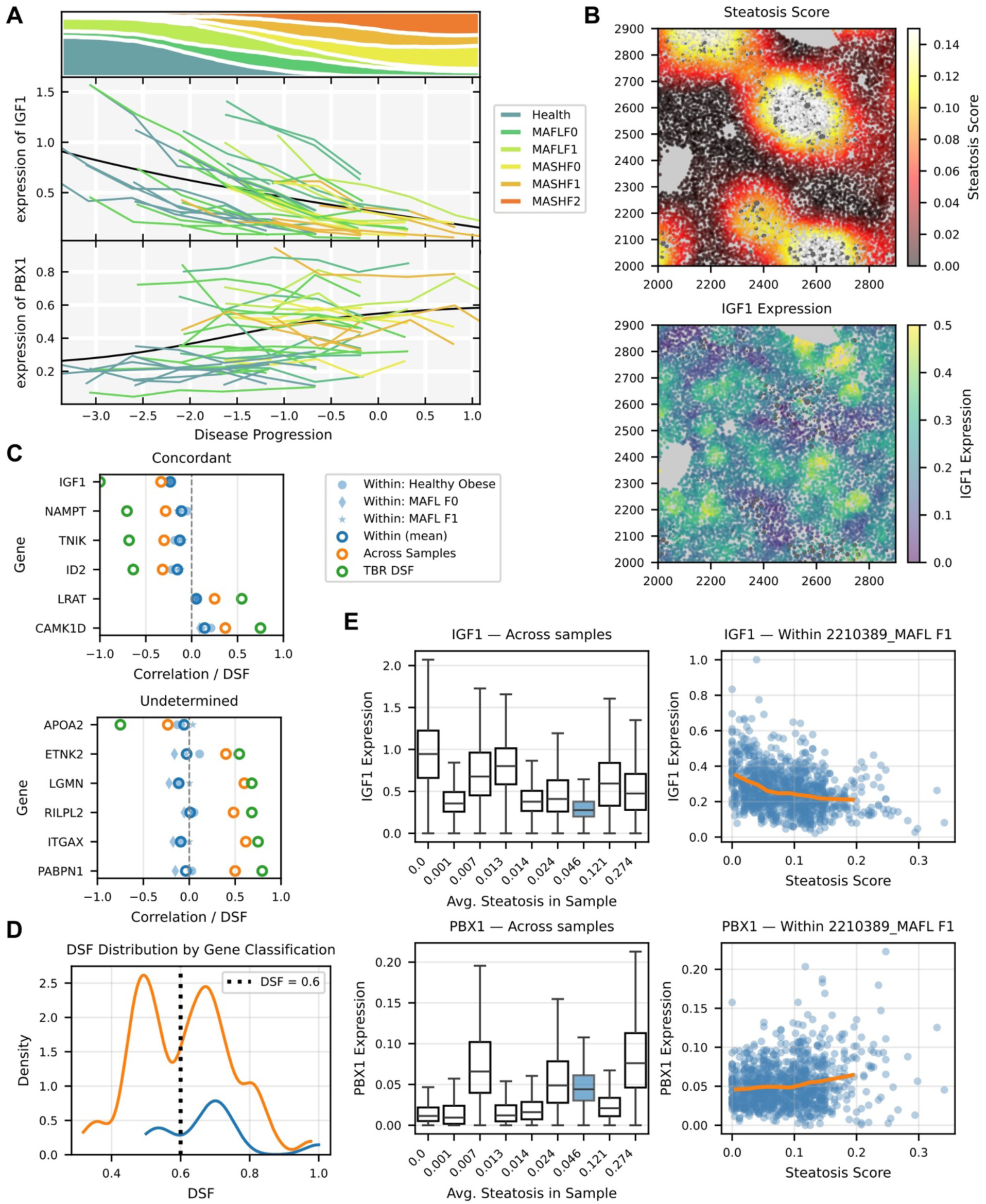
Within-patient heterogeneity and validation with Spatial Tx. **(A)** TBR expression profiles. Examples of two genes, whose overall expression changes with disease progression. IGF1 exhibits within-patient concordance with the overall trajectory (top), while the expression of PBX1 is relatively constant across all cells within each patient (bottom). The overall trajectory is represented by the black line, while the colored lines correspond to individual patient trajectories. The streamgraph at the top shows the proportion of cells from each patient category that were mapped to the corresponding region along the disease continuum. **(B)** Spatial transcriptomics measurements. The top panel shows the spatial map of steatosis scores for individual cells, defined based on their proximity to lipids. The bottom panel displays the corresponding spatial expression map of IGF1. Large gray areas are lipid droplets. Small colored dots are measurements per nuclei. **(C)** Dot plots showing correlation metrics (Within Sample, Across Sample, and DSF) for selected genes categorized as Concordant (top) and Undetermined (bottom). **(D)** Density distribution of DSF scores, illustrating the separation between Concordant (blue) and Undetermined (orange) gene types. The dashed vertical line at 0.6 indicates the proposed threshold DSF value to enrich for Concordant genes**. (E)** Gene expression plotted relative to morphological steatosis. Box plots (left) show the expression of IGF1 (top) and PBX1 (bottom) across all nine samples. The samples are ordered along the x axis based on their average steatosis score, which serves as a patient-level disease label. Scatter plots (right) show the relationship between single-cell gene expression and individual cell steatosis scores within a representative MASLD F1 sample. That sample is highlighted in blue in the boxplot.

To validate that high-DSF genes truly reflect within-tissue cellular disease progression rather than bulk tissue-level variation, we acquired spatial transcriptomics data with matching histopathological stains from nine liver specimens spanning the range of MASLD stages. For each cell, we established a steatosis score by measuring the cell’s proximity to surrounding lipid droplets (Figure 3B). We then isolated all genes that had a significant correlation with disease progression across samples and separated them into “Concordant” and “Undetermined” categories based on whether the correlation also held within the three samples with the largest dynamic range of steatosis (Figure 3C).

We observed that out of the 66 genes correlated with steatosis in our spatial transcriptomics data and expressed at sufficient levels, the proportion of truly concordant genes is relatively small (11 concordant, 55 undetermined). This is likely because the noisy nature of measurements prevents accurate detection of small effect sizes. Nevertheless, the DSF metric helps enrich for the concordant subset (Figure 3D) by establishing an empirical threshold of 0.6 that coincides with the overall bimodal pattern. *PBX1* and other genes below this threshold are almost always in the undetermined category (3 concordant, 25 undetermined), while genes above the threshold are more likely to reliably exhibit intra-patient concordance (Figure 3E). By filtering out negative examples that only exhibit across-sample correlation, the DSF metric helps to identify genuine, mechanistically relevant genes like IGF1 that track with physical disease pathology. Crucially the DSF computation only requires standard single-cell RNAseq data, bypassing the need to run expensive and time-consuming spatial transcriptomics experiments.

### TBRs help enrich for generalizable blood-based biomarkers

Definitive assessment of disease typically requires tissue biopsy, which is invasive, impractical for repeated sampling, and associated with procedural risk. These limitations restrict the ability to detect disease early, monitor its progression, and accurately stage patients. Identifying reliable blood-based biomarkers therefore represents a critical step toward non-invasive disease staging and monitoring.

We examined two publicly available datasets that measure protein abundance in the blood of MASLD/MASH patients using Olink (UK Biobank, n=858; Yang et al.^21^, n=8217). We identified proteins whose levels change significantly with disease in the same direction across both cohorts, yielding 265 candidate biomarkers (Figure 4A). To independently evaluate this initial pool of biomarker candidates, we also collected a confirmation dataset using a different proteomic platform (SomaLogic, n=80) on blood samples acquired from the iSpecimen biobank. Only 42 of the 265 candidates generalized across measurement technologies (Figure 4A), highlighting the challenge of discovering robust blood-based biomarkers from proteomics data alone. We hypothesized that TBR models constructed on single-cell tissue data can help select for proteins that better generalize between independent populations and across measurement technologies, by providing tissue-level explanations for the change in protein levels.

**Figure 4:**
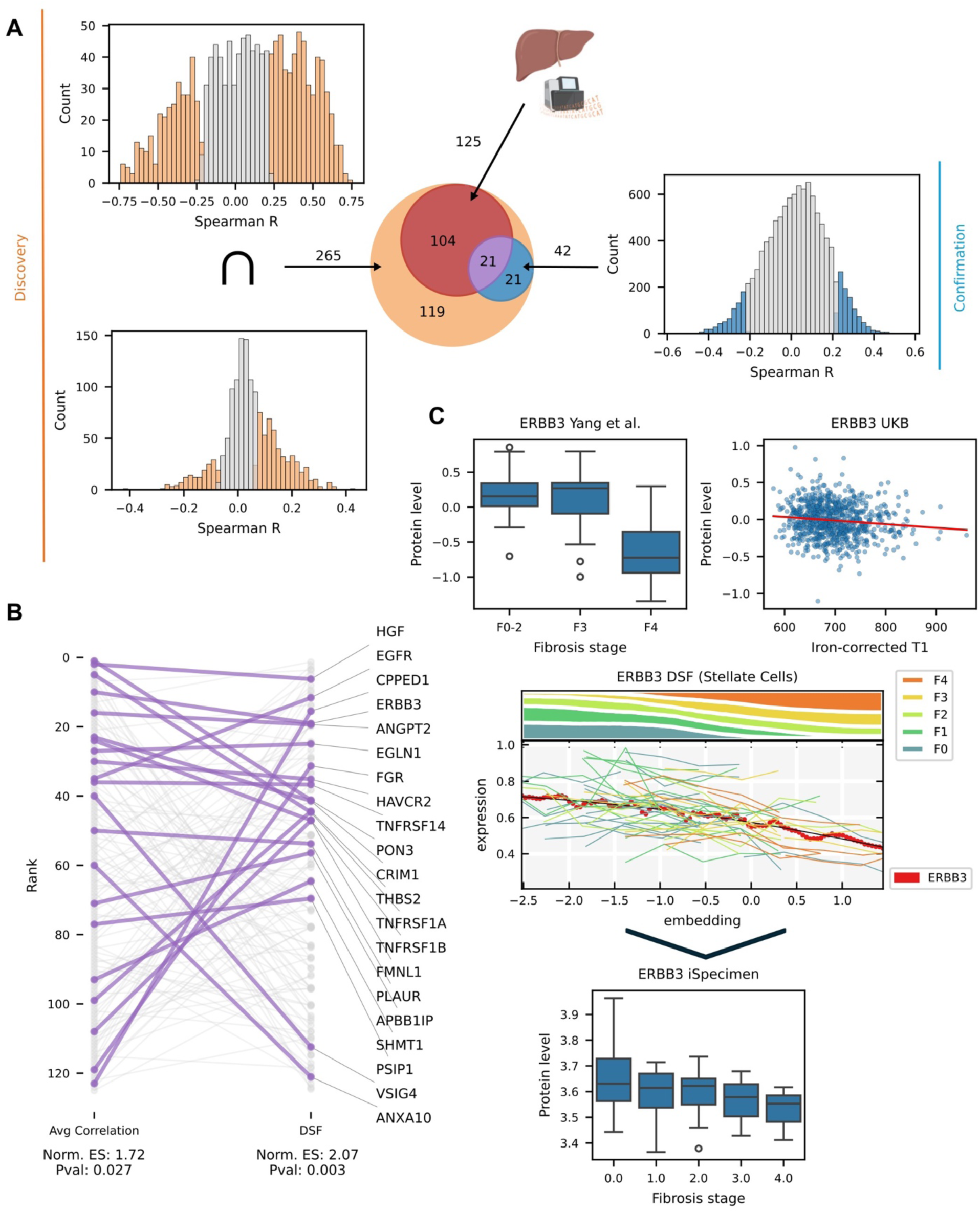
TBR models help identify generalizable blood-based protein biomarkers of liver fibrosis. **(A)** Spearman correlation of protein measurements from two discovery cohorts, UK Biobank (bottom-left) and Yang, et al. (top-left), against fibrosis scores. Proteins showing significant (p < 0.05; Student’s t-distribution) correlations in the same direction with disease progression in both discovery cohorts define the initial pool of candidate biomarkers (orange circle in Venn diagram). The blue circle captures the subset of those proteins that additionally show significant correlation in the same direction in the confirmation set. The red circle represents candidate biomarkers whose genes are also expressed at the mRNA level in single-cell liver data. The purple intersection represents proteins that meet all three criteria. **(B)** Bump chart comparing protein rank across two independent scoring criteria: blood-based correlation (average Spearman statistic across UKB and Yang cohorts) and maximum DSF across liver cell types. Rank 1 indicates the strongest score. Each line represents a protein-coding gene from the initial pool of candidate biomarkers that is also expressed at the mRNA level in the liver tissue (i.e., the red circle from panel A). Of those, protein that also generalize to the confirmation cohort are highlighted in purple with labels; all others are shown in grey. **(C)** ERBB3 is an example protein that had strong correlation with fibrosis in the discovery datasets and high DSF. The same trend generalized the validation dataset. The boxes in boxplots represent the interquartile range (Q1 to Q3) with the line corresponding to the median; the whiskers extend 1.5*IQR from the box edges with individual points highlighting outliers beyond this range. The DSF plot contains the same elements as those in Figure 3A.

Simple tissue-based approaches, such as differential gene expression applied to bulk mRNA measurements or cell type specificity, are not able to identify the initial set of blood-based biomarker candidates (Suppl. Figure 2A, B). This suggests that bulk-level comparisons differ substantially between mRNA and protein levels and a portion of robust biomarker candidates is secreted by tissues other than the liver in response to MASLD, degraded post-transcriptionally, or not secreted into the bloodstream. Without considering tissue measurements, average protein-level correlations with disease severity in the discovery cohort yield a modest but significant enrichment of the robust markers near the top of the ranking (GSEA NES = 1.72 p = 0.027; Figure 4B). Taken together, these initial results suggest that there is room for improvement over blood-based measurements, but that improvement cannot be achieved with bulk-level tissue measurements or cell type specificity alone.

Out of several TBR-derived ranking strategies, signal-to-noise ratio, p-value, or correlation between gene expression profiles and the TBR disease trajectory did not enrich for generalizable biomarkers (Suppl. Figure 2C). Conversely, DSF was consistently able to identify robust, generalizable protein biomarkers (NES = 2.07, p = 0.003; Figure 4B), with 15 of the 21 liver-expressed confirmed biomarkers appearing in the top third of the ranking. The ranking by DSF was complementary to the signal present in protein-based correlations, and combining the two rankings via the geometric mean produced a minor improvement over both (NES = 2.2, p < 0.001). One of the top-ranked genes was ERBB3, which encodes Erb-B2 Receptor Tyrosine Kinase 3 (ErbB3, HER3), a member of the epidermal growth factor receptor (EGFR) family. ERBB3 has been implicated in multiple liver diseases, including liver steatosis^22^. ERBB3 levels were negatively associated with fibrosis progression in all three proteomic datasets, as well as in snRNA-seq data (Figure 4C), suggesting that the proteomics signal is robust and generalizable, and that tissue-based models help select for it.

### TBRs enable temporal analysis of regulatory networks, pathways, and perturbation responses

Beyond gene expression trajectories, TBRs enable two critical temporal capabilities that extend target discovery: (i) latent activity inference for regulatory factors and pathways, and (ii) alignment of cellular perturbation experiments to human disease trajectories. The activity of transcription factors (TF) and receptors is not directly informed by their mRNA expression and must instead be inferred from the expression changes in downstream genes affected by their activity. The Temporal Biodynamics platform performs latent activity inference using a hierarchical regulatory network constructed by integrating three complementary databases: CellPhoneDB^23^ for ligand-receptor interactions, Pathway Commons^24^ for receptor-to-TF signaling cascades, and CollecTRI^25^ for TF-gene regulatory relationships. This multi-layer architecture captures the full signaling cascade from extracellular ligands through membrane receptors and intracellular signaling to transcriptional regulation. By aggregating temporal gradients of downstream gene expression along the disease trajectory, the Temporal Biodynamics platform infers upstream regulator activity through the collective behavior of their well-measured targets, circumventing the challenge of having low mRNA expression of the regulators themselves (Figure 5A).

**Figure 5:**
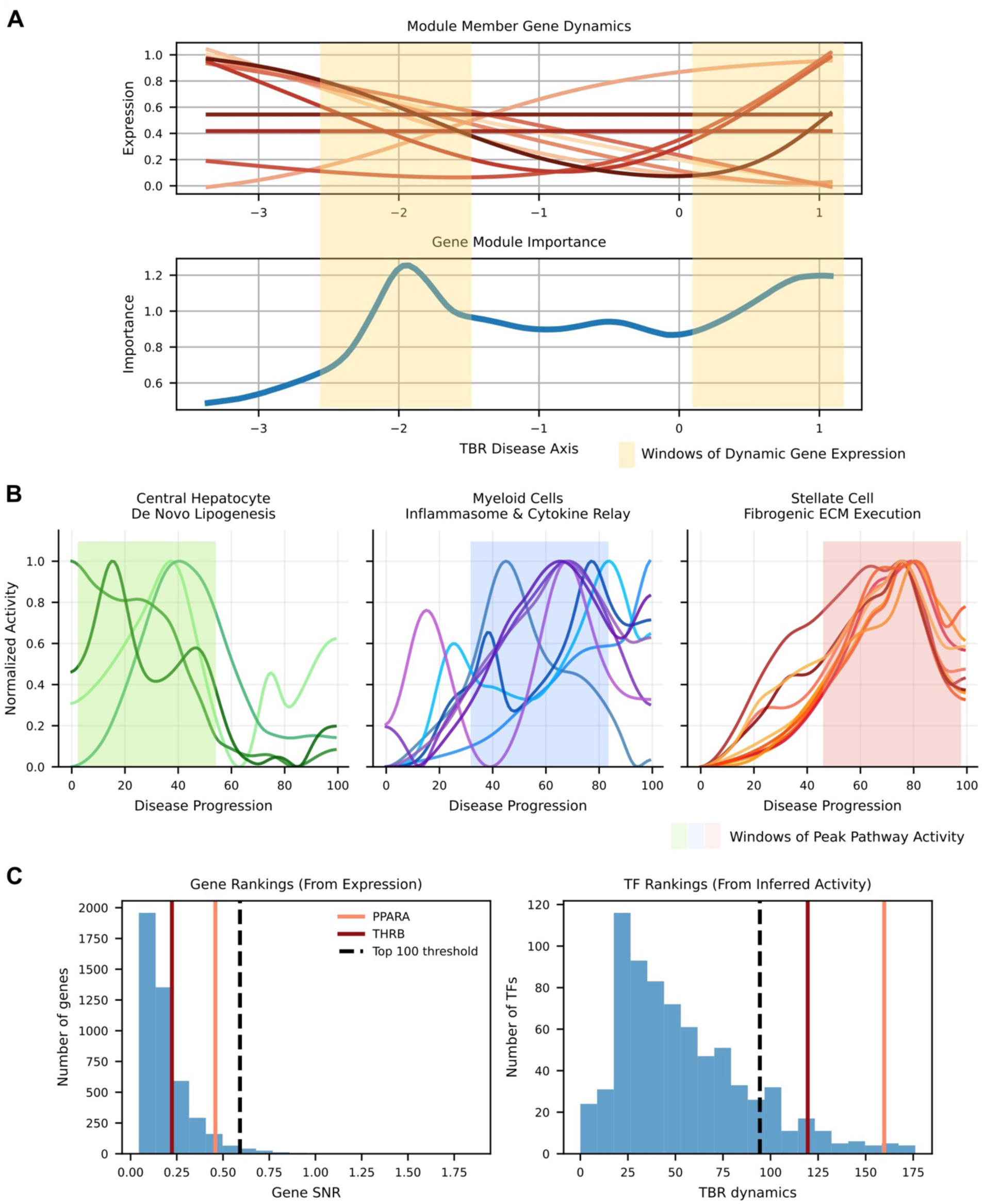
Temporal profiles of gene regulation and pathway activity. **(A)** Importance of a gene module is computed by aggregating temporal gradients of member genes; windows of peak activity correspond to regions where member genes are most dynamic along the TBR disease trajectory. **(B)** Pathway activity inference enables temporal disease understanding. Shown is the lipotoxin relay, a known cascade of disease processes in MASLD. Each panel shows normalized pathway activity trajectories across the inferred disease axis. Each curve represents a pathway related to a particular biological phase. Windows of highest activity are highlighted with color rectangles. **(C)** The distributions of signal-to-noise ratio (SNR) measured in the original mRNA expression space (left) and the inferred transcription factor (TF) activity space (right). The data in both histograms is split into 20 uniform bins. The SNR threshold corresponding to the top 100 genes or TFs is highlighted with a dashed black line. The ranks of PPARA and THRB, known regulators of MASLD progression, are highlighted with colored vertical lines.

The temporal gradient aggregation approach can be generalized beyond the inference of TF activity to characterize temporally resolved importance of any gene group, including those encoding pathway-gene membership information from established databases, such as KEGG^26^ and Reactome^27^. By aggregating temporal gradients of genes belonging to a particular pathway, we computed a measure of that pathway’s activity at each point along the TBR disease trajectory (Figure 5A, B). The resulting activity inference enables temporal reasoning about the cascade of biological processes in disease progression. In MASH, gradient aggregation recapitulates the known lipotoxin relay circuit and its progression through three temporally ordered phases linking hepatocyte lipid stress to fibrotic scar^28,29^ (Figure 5B): in the early phase, central hepatocytes accumulate lipids and induce lipotoxic signaling to neighboring cells; in the mid phase, Kupffer and macrophage cells amplify inflammation and direct cytokines at stellate cells; in the late phase, stellate cells contribute to fibrogenesis, depositing and crosslinking collagen, remodeling the matrix, and committing as myofibroblasts to produce irreversible fibrotic scarring.

Aggregation of mRNA expression gradients also successfully identifies known disease-associated regulators that may be obscured by expression-only analyses. For example, key regulatory proteins PPARA and THRB, known for their role MASLD pathophysiology, are prominently highlighted by latent activity inference despite their low transcriptional expression and dynamics (Figure 5C). Benchmarking against ordinal stage-based inference and established methods (AUCell^30^, VIPER^31^) revealed that TBR’s continuous embedding provides superior resolution for detecting subtle regulatory dynamics, achieving significantly higher enrichment for known disease-associated TFs and receptors (Suppl. Table 2).

### TBRs enable causal inference

In addition to protein activity inference, TBRs also enable contextualization of cellular perturbation experiments within the framework of human disease progression. A fundamental challenge in drug discovery is understanding which aspects of disease are captured by *in vitro* assays and how those aspects are modulated by compound treatments, gene knockdowns, or disease-inducing stimuli in transcriptional space. We expect that perturbations giving rise to disease-modifying therapies reverse the gene expression patterns observed in a TBR model of disease progression. To quantify this, we define a shared disease axis, against which gene knockdowns and other perturbations are subsequently evaluated (Figure 6A).

**Figure 6:**
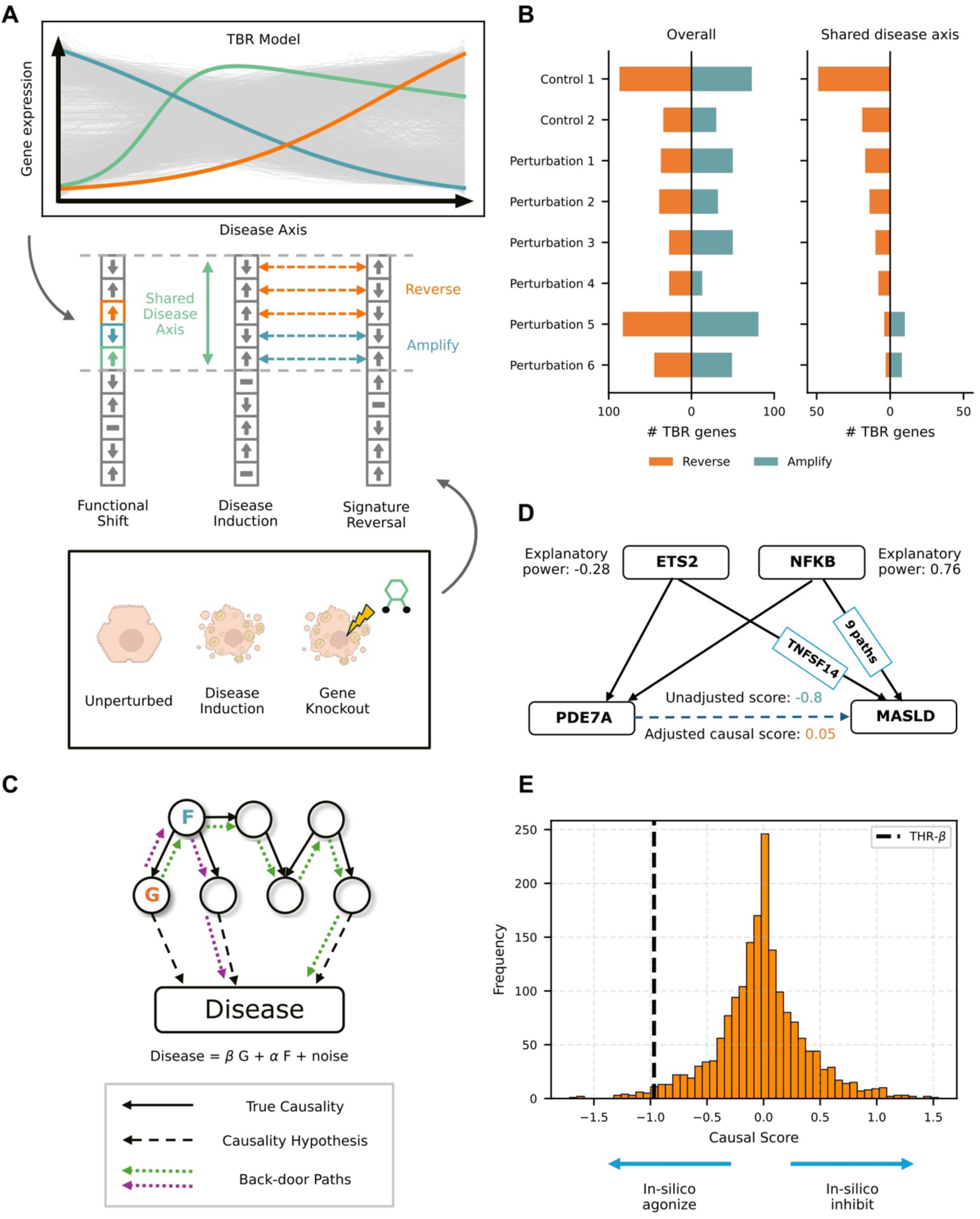
Causal inference over temporal disease models. (**A)** A schematic representation of causal inference using post perturbational data. A shared disease axis is defined as all genes that move in response to disease induction in vitro in the same direction as in a TBR model. Perturbations in individual genes are then evaluated relative to the shared disease axis. **(B)** Gene expression changes from a post-perturbational screen evaluated with and without a shared disease axis. **(C)** A schematic illustration of how causal hypotheses are constructed using predefined gene regulatory networks (solid arrows). The orange node (G) represents a candidate target gene, and the blue node (F) represents a confounder identified in the regulatory network. Dashed arrows represent hypothesized causal relationships between genes and disease based on TBR-derived gene expression profiles. Two alternative backdoor paths (purple and green dotted arrows) connect G to the disease node through the intermediate regulator F. Because both backdoor paths pass through F, conditioning on F alone blocks all confounding paths and enables unbiased estimation of the direct causal effect of G on disease. Each hypothesized causal relationship (dashed edge) is evaluated for rejection based on the resulting estimate and its comparison to the unadjusted equivalent. **(D)** Observed association versus causal effect of PDE7A on MASLD. Although. **(E)** Identification of a clinically validated therapeutic target through causal analysis.

The shared disease axis is constructed by identifying genes whose direction of change is concordant between the TBR-derived human disease trajectory and in response to a stimulus (e.g., a chemical or genetic trigger) applied in an *in vitro* model to recapitulate the disease state in cultured cells. Test perturbations, such as compound treatments or gene knockdowns, are then scored relative to this disease-induction baseline: a gene is counted as reversed if the test perturbation pushes it in the direction opposite to disease induction, and as amplified if it is pushed further along the disease direction. Without first establishing the shared disease axis, comparing test perturbations against the full set of disease-relevant TBR genes leads to ambiguity, where approximately half of measured gene expression changes appear to reverse direction, while the other half appears to be amplified (Figure 6B). This makes it challenging to select the most promising candidates for further follow-up. Restricting the reversal/amplification scoring to the shared disease axis resolves the ambiguity and enables effective prioritization of disease-reversing perturbations (Figure 6B).

A fundamental challenge in target identification is distinguishing genes that causally drive disease progression from those that are merely correlated with disease severity. High correlation between a gene’s expression and disease state may arise through multiple mechanisms: the gene may directly influence disease pathology, it may be a downstream consequence of disease processes, or its apparent association may be mediated entirely through confounding relationships with other genes and proteins. To address this challenge, we developed an approach that utilizes the framework of backdoor paths and adjustment sets from causal inference theory^32^ to provide more explanatory power to the temporal gene expression patterns identified by our TBR models (see Methods).

In this framework, we represent gene regulatory networks as directed acyclic graphs with edges encoding known regulatory relationships between genes based on curated databases. A gene’s observed correlation with disease progression may be explained by “backdoor paths” -indirect paths through the regulatory network that create spurious associations (Figure 6C). For example, a gene may appear to be strongly correlated with disease progression even if it has no direct causal effect, if it is regulated by another protein that also drives disease progression through an alternative path. We evaluate whether a gene’s association with disease persists after controlling for proteins that block all alternative paths and reject the causal hypothesis when a lower association with disease is observed by this adjustment.

For example, pentoxifylline, a non-selective phosphodiesterase (PDE) inhibitor, showed mixed results in small, randomized trials for MASH, with reports of significant improvement in histological NAFLD activity score but not steatosis, hepatocellular ballooning or fibrosis^33,34^. We asked if the causal analysis between the target of pentoxifylline, Phosphodiesterase 7A (PDE7A), and MASLD can help explain the lack of sustained efficacy. We observed that the association of MASLD with PDE7A is largely explainable through alternative non-causal paths in the corresponding regulatory subnetwork (Figure 6D). Specifically, PDE7A’s expression is regulated by two upstream transcription factors, ETS2 and NFKB. ETS2 contributes one alternative regulatory path to the MASLD node through a downstream gene, while NFKB contributes nine such alternative paths, all of which are ancestors of the MASLD node in the causal network. Together, these upstream regulators independently drive disease progression, creating spurious association between PDE7A and MASLD. After adjusting for all confounding paths, the direct causal effect of PDE7A on disease progression is 0.05, which is substantially lower than the unadjusted score of -0.8, computed without considering upstream regulators. Controlling for upstream activity of ETS2 and NFKB reveals that PDE7A is not likely to be a causal driver, providing a potential explanation for the late-stage clinical trial failure.

To systematically evaluate the ability of causal scores to distinguish true therapeutic targets from spurious associations, we applied our adjustment framework at scale to all genes characterized by our TBR model of MASH. We then focused on THRB (thyroid hormone receptor beta), the target of resmetirom, which is the first FDA-approved therapy for MASH. While hundreds of genes significantly correlated with disease progression, only a small subset retained their strong correlation after accounting for confounding regulatory paths. This subset includes THRB (causal score = −0.97), whose association with disease progression cannot be explained by indirect effects through the regulatory network, providing computational evidence for its role as a causal disease driver (Figure 6E).

## Discussion

Confounding signals are ubiquitous in real-world molecular datasets. Failure to properly account for them during target identification can lead to spurious correlations that result in failed clinical trials and wasted resources. Bulk tissue comparisons fail to capture cellular heterogeneity and cannot effectively disentangle disease mechanisms from patient-specific effects. Similarly, existing pseudotime methods have been repeatedly shown to work well in modeling developmental biology but are ill-suited for real-world disease applications where the primary signal of interest is substantially more subtle than cell differentiation. Here, we present a computational framework that models disease as a continuous cellular process and explicitly accounts for patient-to-patient variability, while requiring only cross-sectional sampling commonly found in disease studies. Furthermore, by aligning within-patient cellular heterogeneity to the cohort-level disease trajectory, the “perfect internal control” argument allows for identification of gene expression changes that are truly correlated with disease progression and not driven by unrelated (and possibly unmeasured) confounders.

Target identification efforts do not typically suffer from lack of options. The challenge is rarely in finding new targets, but rather in prioritizing the multitude of nominated genes of interest. Excluding a true negative from the nomination list can be extremely valuable, as it can save a tremendous amount of time and resources spent evaluating an ineffective treatment option in a clinical trial. The DSF and causal score metrics presented in this study provide natural negative filters, allowing researchers to flag nominated gene targets that have spurious correlations to disease progression due to confounding factors or upstream regulators.

Clinical trials may also fail if treatment is given to the wrong patient subpopulation. As we highlighted above, MASLD progression goes through distinct stages of lipogenesis, inflammation and recruitment of stellate cells, followed by fibrogenesis. A treatment targeting lipid accumulation, for example, may not be effective in late-stage fibrosis patients. The temporal models presented here can help avoid incorrect timing of interventions by matching target nominations to specific temporal windows along the disease. Once the window of relevance has been identified, the right segment of the population must be recruited for a clinical trial. While this can be accomplished using electronic health records alone^35^, a stronger biomarker is expected to be blood-based, where the abundance levels of proteins serve as a proxy for the molecular state of the tissue. We have shown that explicitly correlating blood-based protein measurements to the gene expression trends captured by our temporal models can help identify robust biomarkers that generalize even across measurement technologies, where platform-specific effects can otherwise introduce ambiguity^36^.

In chronic diseases, the earliest (“silent”) stages carry subtle signals that escape detection using conventional methods and present a unique opportunity to intervene before a decline in quality of life. Efforts are already underway to help diagnose patients early and include, e.g., voice changes as an early marker of Parkinson’s Disease^37^. Once such detection methods are accepted in clinical practice, the field will be primed for intervention at the critical early stages of chronic conditions. Temporal modeling of disease progression is poised to revolutionize target identification and prioritization, offering insights into early disease processes that are otherwise hidden by traditional case-control comparisons. Furthermore, the Temporal Biodynamics platform presented here demonstrates that early disease characterization is possible in scenarios where longitudinal data is scarce or unavailable, by leveraging cellular heterogeneity of snapshot samples. This is a critical component of early treatment, where one cannot wait until further disease progression to collect longitudinal measurements.

Our study has several limitations, stemming primarily from the reliance of the platform on single-cell sequencing. First, there is no direct connection to genetics. If a particular gene variant is implicated in disease predisposition, that information is invisible to our temporal models, unless it manifests in mRNA expression. Second, disease drivers with low mRNA expression may be inadvertently filtered out by quality control procedures. While we proposed ways to infer protein activity from downstream mRNA patterns, our method relies on known gene regulatory relationships, which may not be readily available for understudied proteins. Lastly, alternative splice forms are sometimes implicated in disease etiologies (e.g., androgen receptor splice variant 7 is an important oncogenic driver in prostate cancer^38^), and the 3’ capture commonly used by single-cell sequencing platforms severely limits the detection power of such variants. Overall, these limitations are compensated for by the wide availability of single-cell and single-nuclei sequencing datasets, and the ability of our platform to make use of within-patient cell heterogeneity without an explicit access to spatial information.

## Methods

### Overview

The TBR computational framework models chronic disease progression as a continuous cellular process by learning a one-dimensional embedding that arranges individual cells along a disease trajectory (though multi-dimensional are compatible with the framework). The method addresses fundamental challenges in chronic disease modeling from single-cell RNA sequencing data, including cross-sectional sampling constraints where each patient provides only a snapshot of cellular states, substantial within-patient cellular heterogeneity that is obscured by discrete clinical staging, and patient-level biological and technical confounders that mask disease-specific signals.

### Quality Control

Single-cell RNA (snRNA-seq) sequencing data was processed using standard quality control procedures. For each snRNA-seq dataset considered in the study, cells were filtered based on minimum gene counts (typically >200 genes per cell), median absolute deviation (MAD) cutoffs for total counts (5 MADs) and mitochondrial content (5 MADs from median, with absolute maximum of 20%), and doublet removal using Scrublet^39^ with expected doublet rates calculated per sample (0.8% per 1000 cells for 10X Genomics data). Gene expression matrices were normalized using library size normalization and log-transformation (log(counts per 10,000 + 1)). Cell type annotation was performed using established marker genes and validated against reference atlases.

### Feature Selection and Pathway-Constrained Dimensionality Reduction

To identify disease-relevant genes while maintaining biological interpretability, we projected the original gene expression values into a pathway-constrained latent space using PathwayAE, a specialized sparse autoencoder architecture where each latent dimension corresponds to a known biological pathway. The encoder maps gene expression to pathway activity scores, such that each latent dimension aggregates only genes belonging to the corresponding pathway. The decoder is a dense network. Pathway definitions were taken from publicly available databases include KEGG^26^, Reactome^27^, and MSigDB^40^.

After training, the PathwayAE embeddings were filtered to remove latent dimensions more strongly associated with sample-level variation than disease progression, reducing technical and biological confounding. For each latent dimension *p*, we computed F-statistics quantifying its association with disease stage and relevant sample attributes. Latent dimensions were ranked by the differences between F values, and the optimal number of dimensions to retain was determined using the kneedle algorithm^41^. All other dimensions were dropped.

### Ordinal Regression Neural Network for Continuous Embedding

The ordinal module of the TBR framework maps each cell’s pathway-level representation to a continuous disease coordinate that reflects its position along a shared disease progression continuum. Starting from the pathway-constrained latent embedding of each cell, the ordinal module learns a scalar coordinate, whose larger values correspond to more advanced disease states. Crucially, the disease coordinate is not directly supervised. Instead, it emerges as a latent variable from an ordinal regression task trained using patient-level clinical stage labels. This formulation enforces monotonic consistency with clinical staging while preserving transcriptional continuity, enabling recovery of a continuous disease trajectory from discrete supervision. We utilized either the CORAL^42^ or the CORN^43^ neural network architecture, depending on the dataset. In general, we observed that CORAL was better suited to datasets with a smaller number of patient samples, due to its constraint that all disease stage transitions share a common weight vector. To ensure smooth transitions between neighboring states, CORN-based models were additionally regularized using gradient penalties^44^. To avoid overfitting, all models were trained with early stopping by evaluating performance on a withheld subset of data.

### Gene Expression Profile Construction

To construct smooth gene expression profiles along the learned disease severity embedding, we applied kernel-weighted averaging at regularly spaced coordinates. For each of *M* = 100 positions *x_m_* along the embedding axis (linearly spaced between the 1st and 99th percentiles of cell embeddings), we defined a kernel *K_M_* that assigns weights to nearby cells:

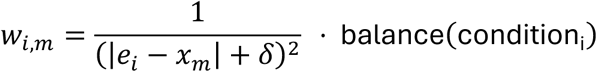

where 𝑒_#_ is the embedding position of cell 𝑖, 𝛿 = 0.5 is a distance offset to prevent division by zero, and balance(*condition_i_*) upweights cells from underrepresented disease stages to ensure equal contribution across conditions. Weights are row-normalized as *ȃw _i,m_* = *w_i,m_*/ ∑*_j_w_i,m_*, and the kernel is restricted to the 𝑁 = 2,000 nearest cells to each position.

To obtain robust trajectory estimates and quantify their uncertainty, we applied a non-parametric bootstrap at the patient level. In each of 𝐵 = 100 replicates, patients were resampled with replacement, and cell weights were scaled by the number of times each cell’s patient was drawn before renormalization:

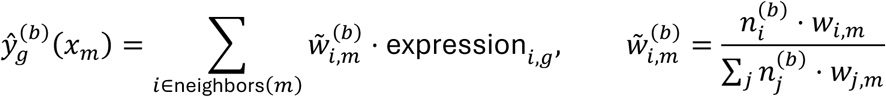

where 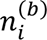 is the number of times cell *i*’s patient was drawn in replicate 𝑏. The reported mean trajectory and standard error are then the mean and sample standard deviation across replicates:

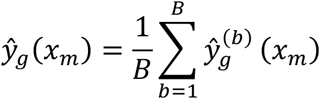

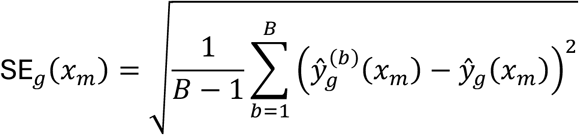

The standard error captures variability in the smoothed trajectory attributable to donor-level sampling, providing a coordinate-wise measure of confidence in the estimated expression profile. The resulting output consists of expression trajectories 𝑦H_’_(𝑥) and standard errors 𝑆𝐸_’_(𝑥) for each gene 𝑔 across the embedding axis.

### Signal Robustness and Statistical Significance

Given a gene expression profile, a signal-to-noise ratio (SNR) quantifies overall trajectory dynamics relative to local variability:

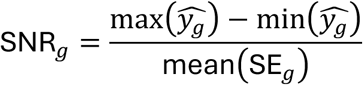

where the signal is the total range of expression change along the trajectory and noise is the average standard error across trajectory positions.

### Statistical significance testing

To assess whether observed gene dynamics reflect true disease progression rather than random variation, we generated null distributions by training parallel TBR models with shuffled patient labels. For each gene, we computed the empirical p-value as the fraction of null models with SNR greater than or equal to the observed SNR. P-values were adjusted for multiple testing using the Benjamini-Hochberg procedure (FDR < 0.05).

### Directionally Similar Fraction (DSF)

To identify genes that track with cellular disease progression within individual patients (rather than varying only at the bulk tissue level), we computed the DSF metric. For each gene g and patient p, we determined whether the gene’s expression change within that patient’s cells (along the embedding) matched the population-level trend direction. DSF is the fraction of patients showing concordant within-patient directionality:

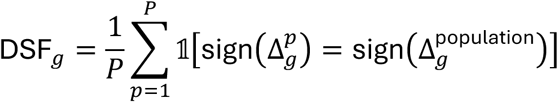

where Δ_g^p is the expression change within patient p’s cells and Δ_g^population is the overall population trend. Genes with DSF > 0.6 were classified as having high concordance with the overall disease trajectory.

### Validation Dataset and Ground Truth Construction

To rigorously evaluate TBR performance under controlled conditions with known ground truth, we constructed a disease dataset from the mouse embryonic fibroblast (MEF) to induced endoderm progenitor (iEP) reprogramming time series^12^. The original dataset was downloaded from Gene Expression Omnibus (accession number GSE101099) and contains ∼84,000 cells sampled at 8 time points (days 0, 3, 6, 9, 12, 15, 21, 28) during transcription factor-mediated reprogramming.

### True cell-level scoring

We computed continuous iEP identity scores for each cell using quadratic programming, following the approach described in the original paper^12^. For each cell, we solved the constrained least-squares problem: minimize ||Ax - b||² subject to x ≥ 0 and Σx = 1, where A = [MEF_profile, iEP_profile] is a matrix of reference expression profiles and b is the cell’s expression vector. The MEF reference profile was extracted from day 0 cells, while the iEP reference profile was obtained by clustering day 28 cells (Leiden algorithm, n_neighbors=15, resolution=0.5) and selecting the cluster with highest expression of endoderm markers (Apoa1, Cdh1, Hnf4a, Foxa2, Gata4). The optimization was performed using 1000 highly variable genes (Seurat v3 method). The resulting iEP score—the coefficient for the iEP reference profile—provides a continuous measure of differentiation state ranging from 0 (MEF-like) to 1 (iEP-like), serving as ground-truth cell-level disease labels.

### Disease stage discretization

We discretized the continuous iEP scores into 6 disease stages using quantile-based binning: Stage 0 (iEP score < 0.167, “healthy”), Stage 1 (0.167-0.333), Stage 2 (0.333-0.5), Stage 3 (0.5-0.667), Stage 4 (0.667-0.833), and Stage 5 (≥0.833, “diseased”). This discretization mimics clinical staging systems while preserving the underlying continuous cellular distribution.

### Simulated patient generation

We generated 30 synthetic patient samples, each containing ∼2000 cells, to simulate a cross-sectional disease cohort. To create realistic heterogeneous cellular mixtures, we first binned all cells into 100 equally-spaced bins based on their continuous iEP scores. Each patient was assigned a discrete disease condition label (0-5) corresponding to the six disease stages defined above. For a patient with condition label n, we randomly selected a center bin from the iEP score range corresponding to stage n (e.g., condition 3 patients sample from bins 50-66, corresponding to iEP scores 0.5-0.667), then sampled 2000 cells according to a Gaussian distribution centered at that bin with standard deviation of σ bins. This approach ensures that each patient’s cellular composition is centered within their assigned disease stage while containing cells from adjacent stages, faithfully recapitulating the within-patient heterogeneity observed in real disease cohorts where individual patients at the same clinical stage exhibit varying cellular distributions along the disease continuum.

### Confounder and covariate addition

To recapitulate real-world challenges, we added both technical confounders and biological covariates. For technical batch effects, we identified 7,500 highly variable genes from the original normalized data and randomly selected 30% of these genes to be affected by sample-specific noise in each patient. We applied multiplicative noise (lognormal with σ = 0.2) and additive noise (Gaussian with σ = 0.1 × mean expression) to the selected genes in raw count space. For biological covariates, we assigned each patient a random age (uniformly sampled from 20-80 years) and sex (randomly assigned as male or female). Covariate effects were applied to gene expression by modulating a subset of genes based on these patient-level attributes: age effects were modeled as linear trends where gene expression increased or decreased proportionally with age, while sex effects introduced binary shifts in expression levels between male and female samples. The strength of these covariate effects was controlled by a parameter set to 0.5, determining the magnitude of expression changes relative to the baseline disease signal.

### Baseline Methods for Comparison

#### Unsupervised Trajectory Inference Methods

**PAGA (Partition-based Graph Abstraction)** was implemented using Scanpy v1.9.1 with default parameters. Diffusion pseudotime (DPT) is computed along the principal trajectory identified by PAGA.

**Palantir** was implemented using Palantir v1.0.1. Early-stage cells are selected as starting point, and pseudotime is computed using diffusion maps with n_components = 10.

**Diffusion Pseudotime (DPT)** was computed using Scanpy’s DPT implementation with root cell selected from the earliest disease stage.

#### Supervised Label Refinement Methods

**HiDDEN (Hierarchical Differential Expression Network)** was applied to refine discrete stage labels using local neighborhood information in PCA space (k = 30 neighbors).

**MELD (Manifold Enhancement of Latent Dimensions)** was used to compute continuous disease likelihood scores for each cell based on stage label propagation on k-nearest neighbor graph (k = 30).

**Milo (Differential abundance testing)** was used to perform differential abundance testing across disease stages with default parameters. Genes are ranked by association with differentially abundant neighborhoods.

#### Traditional Differential Expression Methods

**PseudobulkDESeq2** aggregates cells to sample-level pseudobulk count profiles per disease stage, then ranks genes by the maximum |Wald statistic| across stage-vs-stage-0 contrasts.

**ANOVA** ranks genes by a one-way F-statistic computed per gene across ordinal disease stage.

**Wilcoxon rank-sum test** is applied in a pairwise fashion between consecutive disease stages. The final gene ranking is given by the minimum p-value across comparisons.

For all methods, we used the same input data (normalized expression matrices, cell type annotations, disease stage labels) to ensure fair comparison. Gene rankings from each method were evaluated using the same evaluation metrics.

### Evaluation Metrics

#### Trajectory Reconstruction Accuracy

For synthetic data with ground-truth cell-level labels t_i:

**Pearson correlation:** r = corr(embedding_i, t_i) measures linear agreement between inferred embedding and true labels.

#### Gene Set Enrichment

To evaluate gene rankings produced by each method, we first defined two sets of ground-truth disease sets:

**Literature-curated disease genes** taken from clinical literature, genetic studies, and pathway databases representing established disease mechanisms for MASLD, including lipid metabolism, inflammatory cytokines, fibrosis markers, insulin signaling components, and others.

**Clinical trial targets** were taken to be genes encoding proteins targeted by drugs that entered a Phase III clinical trial for MASLD or MASH. Data was obtained from ClinicalTrials.gov and DrugBank.

For each method’s gene ranking, we then computed enrichment scores of ground-truth gene sets using the following metrics.

**Normalized Enrichment Score (NES)** is computed using GSEA (Gene Set Enrichment Analysis) algorithm, comparing ranked gene list against ground-truth disease gene sets. Statistical significance is assessed by permutation testing (1000 permutations).

**Precision at k** is the fraction of top-k ranked genes that are in the ground-truth set, for k = 50, 100, 200, 500.

**Area Under Precision-Recall Curve (AUPRC)** summarizes precision-recall tradeoff across all ranking thresholds.

#### Clinical Trial Target Enrichment

For genes targeted by drugs in clinical trials, we computed the following metrics.

**Phase progression score:** Mean rank percentile of genes targeted by drugs reaching each phase (I, II, III, approved), with lower ranks (higher priority) indicating better enrichment.

**Rank-phase correlation:** Spearman correlation between gene rank and maximum trial phase reached, testing whether higher-ranked genes are more likely to reach advanced trials.

#### Transcriptional Continuity

To quantify smoothness of cell ordering:

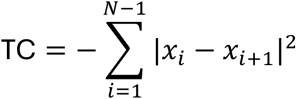

where cells are ordered by embedding position and x_i is the gene expression vector for cell i. Higher (less negative) values indicate smoother transcriptional transitions.

### Single-cell data acquisition for MASLD

Human liver single-nucleus RNA-seq data was acquired from 44 patients spanning healthy, steatosis, MASH, and fibrosis stages. Clinical staging was based on histopathological assessment (NAS score, fibrosis stage).

#### Tissue Procurement

Flash-frozen human liver tissue samples were obtained from Vanderbilt University Medical Center (VUMC) and LifeNET Health. All tissue was stored at −80°C until use.

#### Buffer Preparation

All buffers were prepared fresh on the day of the experiment and maintained on ice throughout. Two-times concentrated salt-Tris (2X ST) buffer was prepared as described in Table 1. Working TST buffer (1X ST supplemented with 0.03% Tween-20 and 0.01% BSA) was prepared as described in Table 2. Protector RNase Inhibitor (Sigma-Aldrich, cat. no. 3335402001; 40 U/µL stock) was added to all buffers to a final concentration of 0.2 U/µL (5 µL per 1 mL buffer). A 1X ST working solution was prepared by diluting 2X ST stock 1:1 with nuclease-free water, supplemented with RNase inhibitor as above.

**Table 1.** Working TST Buffer Composition (per 5 mL)

| Component | Final Concentration |
| --- | --- |
| NaCl | 292 mM |
| Tris-HCl (pH 7.5) | 20 mM |
| CaCl<sub>2</sub> | 2 mM |
| MgCl<sub>2</sub> | 42 mM |
| Nuclease-free water | — |

**Table 2.** 2X Salt-Tris (ST) Buffer Composition.

| Component | Final Concentration |
| --- | --- |
| 2X ST buffer | 1X |
| Tween-20 | 0.03% |
| BSA | 0.01% |
| Nuclease-free water | — |
| RNase inhibitor (Protector, Sigma-Aldrich 3335402001) | 0.2 U/μL |

#### Nuclei Isolation from Frozen Liver Tissue

Approximately 150 mg of flash-frozen liver tissue was weighed and placed in a 1.5 mL microcentrifuge tube on ice. 1 mL of TST buffer was added directly to the tissue. Mechanical dissociation was performed using a pre-chilled disposable plastic pestle with manual grinding until no visible tissue fragments remained (approximately 1–2 minutes). The homogenate was passed through a 40 µm cell strainer and the strainer was rinsed with an additional 1 mL of TST. The filtrate volume was brought to 5 mL with 3 mL of 1X ST buffer. Nuclei were pelleted by centrifugation at 500 × g for 5 minutes at 4°C. Pellet was rinsed gently with 1 mL of 1X PBS followed by resuspension in 1 mL of 1X PBS containing 1% BSA and RNase inhibitor.

#### Flow Cytometric Sorting of Single Nuclei

Propidium iodide (PI) was added to the nuclear suspension at 1 µL per sample and nuclei were sorted by fluorescence-activated cell sorting (FACS) using the 488 nm laser. Nuclei were then counted on a countess II.

#### Single-Nucleus RNA Sequencing

Up to 10,000 sorted nuclei were loaded per channel onto a 10x Genomics Chromium Chip according to the manufacturer’s instructions. Libraries were made using 10x genomics protocol for 3’ gene expression v3.1 with dual index library following manufacturer’s instructions. Libraries were pooled and sequenced on Illumina Nextseq2000.

### Validation of Directionally Similar Fraction

To validate that high-DSF genes truly reflect within-tissue cellular disease progression, we used spatial transcriptomics data from MASLD liver tissues (10X Genomics Xenium platform). For each cell/spot, we quantified local disease severity using imaging-based lipid droplet detection:

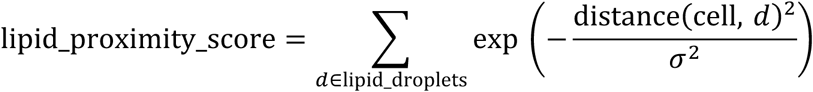

where σ = 50 μm defines the spatial scale. We then tested whether genes with high DSF showed significant within-patient correlation between expression in the spatial transcriptomics data and lipid proximity score, using Spearman correlation computed separately for each patient. Genes with mean within-patient correlation > 0.3 across patients were classified as spatially validated cellular disease genes.

### Biomarker identification

The UK Biobank cohort comprised 858 plasma samples profiled using the Olink Explore 1536 panel, yielding measurements for 1463 proteins. As METAVIR fibrosis scores were unavailable for this cohort, liver iron-corrected T1 (cT1) was used as the disease label^45^ . Spearman correlation of protein levels with cT1 and significance threshold of p < 0.05 were used to identify proteins that are associated with fibrosis severity.

The Yang et al. dataset comprised 82 plasma samples from MASLD patients with histologically characterized fibrosis, profiled using Olink Explore 1536 Proximity Extension Assay. Of those, 26 samples were F0–F2, 14 were F3, and 42 were F4 on the METAVIR scale. In total, 1463 proteins were detected, of which 883 correlated significantly (p < 0.05) with fibrosis stage.

The in-house proteomics dataset was collected using 80 iSpecimen serum samples. Of those, 38 samples were F0, 14 samples were F1, 22 samples were F2, 3 samples were F3 and 3 samples were F4 fibrosis score on the METAVIR scale. Proteomic profiling was conducted by SomaLogic (Boulder, CO, USA) using the SomaScan v11 platform (SomaScan 11K), a highly multiplexed aptamer-based assay that quantifies the relative abundance of 11,083 human protein analytes across a wide dynamic range. Sample processing, hybridization, and signal detection were performed by SomaLogic in accordance with standardized operating procedures. Raw RFU data were processed using SomaLogic’s standard normalization pipeline. In total, 10775 proteins passed all QC metrics, out of which 1666 were significantly (p < 0.05) correlated with fibrosis stage.

Single-cell RNA-seq data were aggregated into pseudobulk profiles prior to differential expression analysis. Cells were grouped by sample identity and gene expression values were summarized within each group using the arithmetic mean across all cells. Differential gene expression between fibrosis stage F4 and F0 was assessed using DESeq2^46^, implemented via the PyDESeq2 package. Fibrosis stage was modeled as the design factor, and size factor estimation, dispersion estimation, and GLM fitting were performed using default DESeq2 parameters with Cook’s distance refitting disabled. Genes were considered to be significantly differentially expression if the adjusted p-values was below 0.05.

Liver-specific genes were identified using RNA expression data from the Human Protein Atlas (HPA) consensus tissue gene expression dataset. For each gene, tissue specificity was quantified using the tau score, defined as τ = Σ(1 −^x ᵢ) / (N − 1), wherêx ᵢ is the normalized expression in tissue i relative to the maximum across all tissues and N is the number of tissues assayed. To identify the set of tissues in which a highly specific gene (τ ≥ 0.95) is enriched, we applied the extended tau method^47^, retaining tissues whose nTPM exceeded the maximum expression minus 1.96 standard deviations of the non-zero expression distribution. Genes with τ ≥ 0.95 and enrichment in liver tissue were retained as liver-specific candidates.

For comparison against tissue-informed biomarker selection, proteins were first filtered to those having sufficient expression at the mRNA level in at least one cell type of relevance (portal hepatocytes, central hepatocytes, macrophages, hepatic stellate cells, Kupffer cells, and cholangiocytes). For this subset, Spearman correlation coefficients with fibrosis severity were averaged across the UKB and Yang cohorts, and the absolute value of the average correlation was used as the pre-ranked metric. Gene Set Enrichment Analysis (GSEA) was then performed using gseapy.prerank against the proteins whose correlation with fibrosis was statistically significant and directionally concordant across all three cohorts (UKB, Yang, and iSpecimen).

For each protein also represented at the mRNA level in the tissue, a global correlation score was computed as the Pearson correlation between gene expression and the disease severity embedding produced by a TBR model. The maximum absolute global correlation score across the six hepatic cell types (portal hepatocytes, central hepatocytes, macrophages, hepatic stellate cells, Kupffer cells, and cholangiocytes) was used as the pre-ranked metric for GSEA against the robust gene set. Notably, no directionality constraint was imposed between the transcriptomic and proteomic signals, as circulating protein levels often do not mirror tissue-level transcription. Similarly, the maximum direction-same fraction (DSF) across six hepatic cell types was used as the pre-ranked metric for GSEA against the robust gene set.

To integrate transcriptomic and proteomic evidence, a combined ranking metric was computed as the geometric mean of the average absolute Spearman correlation across the UKB and Yang cohorts and the maximum absolute DSF score across the six hepatic cell types. The geometric mean was chosen to ensure that proteins scoring highly on both modalities were prioritized, penalizing candidates with strong signal in only one.

### Latent Activity Inference for Regulatory Factors and Pathways

We developed a gradient-based method that infers latent activity of transcription factors (TFs) and receptors from the TBR trajectories of their downstream genes. The approach leverages hierarchical regulatory networks integrating ligand-receptor interactions from CellPhoneDB v4.0^23^, receptor-to-TF signaling cascades from Pathway Commons v12^24,48^, and TF-target gene regulatory relationships from CollecTRI^25^.

For each TF or receptor **r** with known downstream target genes **T_r**, we computed temporal gradients of the TBR trajectories of their targets along the disease trajectory

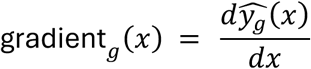

estimated using local linear regression in sliding windows (window width = 10). The latent activity of regulator **r** at position **x** is:

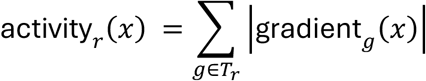

This aggregates the collective dynamic behavior of downstream targets, providing a proxy for upstream regulator activity. We applied Savitzky-Golay filtering (window = 10, polynomial order = 3) to smooth trajectories before gradient computation. Regulators were ranked by maximum activity along the trajectory and validated against known disease-associated factors from literature.

We applied a similar procedure to obtain the activity of predefined sets of pathways. Pathway gene membership information was obtained from existing pathway databases. For each pathway, the temporal gradients of all genes belonging to that pathway were aggregated as above to obtain the overall ‘pathway-activity’.

### Perturbation experiment alignment and scoring

To contextualize *in vitro* perturbation experiments within the human disease trajectory, we first applied DESeq2 to identify differentially expressed genes in each perturbation. Disease-inducing stimuli were contrasted to the corresponding unperturbed state, while compound treatments and gene knockdowns were compared to disease-induced cell states.

Given a set of differentially expressed genes 𝐺_4#BCDBC_from a disease-inducing perturbation experiment, we first establish a shared disease axis with our temporal model by converting the output from DESeq2 into a trinary representation, assigning each gene a value of +1 (upregulated), −1 (downregulated), or 0 (unchanged) based on log2 fold-change and adjusted p-value thresholds (default: |𝑙𝑜𝑔2𝐹𝐶| > 0.5, 𝑝_D4&_ < 0.05). Similarly, we convert gene expression profiles in our temporal models of human disease into a trinary representation based on whether the overall profile goes up, down, or is undetermined due to having low signal-to-noise ratio. The shared disease axis 𝐺_BED?C4_ is defined by the subset of 𝐺_4#BCDBC_ whose trinary representation matches the temporal models.

After establishing a shared disease axis, all other perturbations are scored by counting the number of genes that reverse or amplify the trinary direction in 𝐺_BED?C4_.

### Causal inference

The causal inference framework is grounded in causal graphical models^32^. We represent prior biological knowledge as a directed acyclic graph (DAG), where nodes correspond to genes, transcription factors, receptors, and signaling proteins, and directed edges encode regulatory relationships. The DAG is constructed using the same databases as in protein activity reference. Ligand and target gene mRNAs are treated as observed nodes in the DAG, measured directly by scRNA-seq; receptor and TF proteins are treated as latent nodes whose activities are inferred (see Methods: Latent Activity Inference for Regulatory Factors). A hypothetical *Disease* node is introduced into the DAG to represent disease progression. For a given cell, the value of this node is taken to be the coordinate of that cell along the disease embedding axis, as assigned by our temporal models.

Given a gene regulatory DAG, our goal is to estimate the causal effect of a candidate upstream gene X on *Disease*. Because the true set of causal drivers is unknown a priori, candidate mRNA nodes are nominated from TBR rankings: the top N genes by signal-to-noise ratio (SNR) with a nominal p-value < 0.1 are each connected to the *Disease* node by a hypothetical directed dashed edge (Figure 6c), provided the gene has at least one parent in the causal network (i.e., is embedded in known upstream regulatory biology). Each such edge represents the hypothesis that the gene has a direct causal effect on disease progression. After computing causal scores (see below), genes whose causal score is substantially diminished relative to their marginal association with disease are deprioritized, and their hypothetical edges are removed from the graph; genes that retain a high causal score after adjustment are retained as high-confidence targets.

A key challenge in estimating the causal effect of 𝑋 on *Disease* is the presence of backdoor paths, which induce non-causal associations between 𝑋 and *Disease*. A backdoor path is any path between 𝑋 and *Disease* that begins with an arrow pointing into 𝑋. Figure 6c shows two backdoor paths between 𝑋 and *Disease* represented as dotted purple and green arrows. Such paths create non-causal statistical associations: if a gene 𝑋 is regulated by an upstream transcription factor *Y* that independently drives disease progression, *X* will appear correlated with disease even if it has no direct causal effect.

To recover causal effects, we must block all backdoor paths. Intuitively, *blocking a path* means conditioning on variables that prevent information from flowing along that path. Formally, this is achieved using an adjustment set 𝑆, a set of variables such that conditioning on 𝑆 blocks all backdoor paths between 𝑋 and *Disease* while leaving causal paths intact. In Figure 6c, 𝑆 = 𝑌 blocks both the purple and the green backdoor paths.

To identify valid adjustment sets, we apply a greedy minimization algorithm that starts with the full set of ancestors of *X* and *Disease* restricted to the observed nodes and interactively attempts to remove one node at a time, verifying that the remaining set still blocks all backdoor paths. The algorithm terminates when removing any additional nodes is no longer possible under these constraints.

Given a valid adjustment set 𝑺 = {𝑆_1_, 𝑆_2_ …, 𝑆*_k_*}, we estimate the causal effect of 𝑋 on 𝐷𝑖𝑠𝑒𝑎𝑠𝑒 using ordinary least squares (OLS) linear regression:

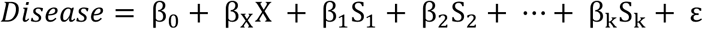

where 𝐷𝑖𝑠𝑒𝑎𝑠𝑒 is the continuous disease score from the TBR embedding, 𝑋is the expression of the candidate target gene, 𝑆_1_, 𝑆_2_ …, 𝑆*_k_* are the expression values (or inferred activities) of the adjustment set variables, and ε is the residual error. The coefficient 𝛽_J_ represents the estimated direct average causal effect of 𝑋on disease after adjusting for all confounding paths through the regulatory network. We refer to 𝛽_J_ as the causal score of gene 𝑋 on *Disease*. If 𝛽_J_ is significantly smaller than the unadjusted correlation of *X* with *Disease*, then we reject the hypothesis that *X* is causal of disease progression. The significance is assessed by comparing causal scores against the background distributions of correlation values and OLS coeáicients across all genes.

