## Supplemental Figures for "Temporal Biodynamics: An AI Platform for Identification of Stage-Relevant Targets and Biomarkers"

### Supplementary Section

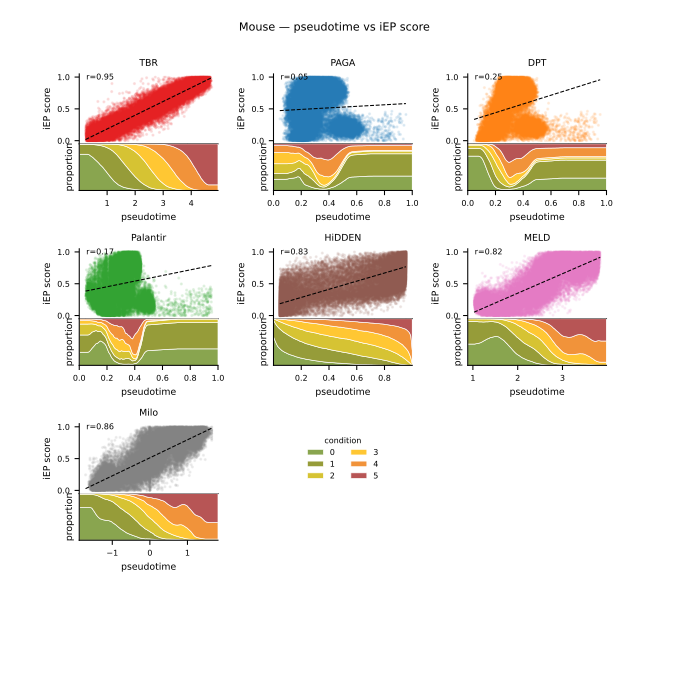

Supplementary Figure 1: **(top)** **Predicted cell-level pseudotime versus true disease (iEP) score for the mouse reprogramming dataset, (bottom) An alternate representation showing proportion plots of discrete disease labels created by binning continuous disease scores.**

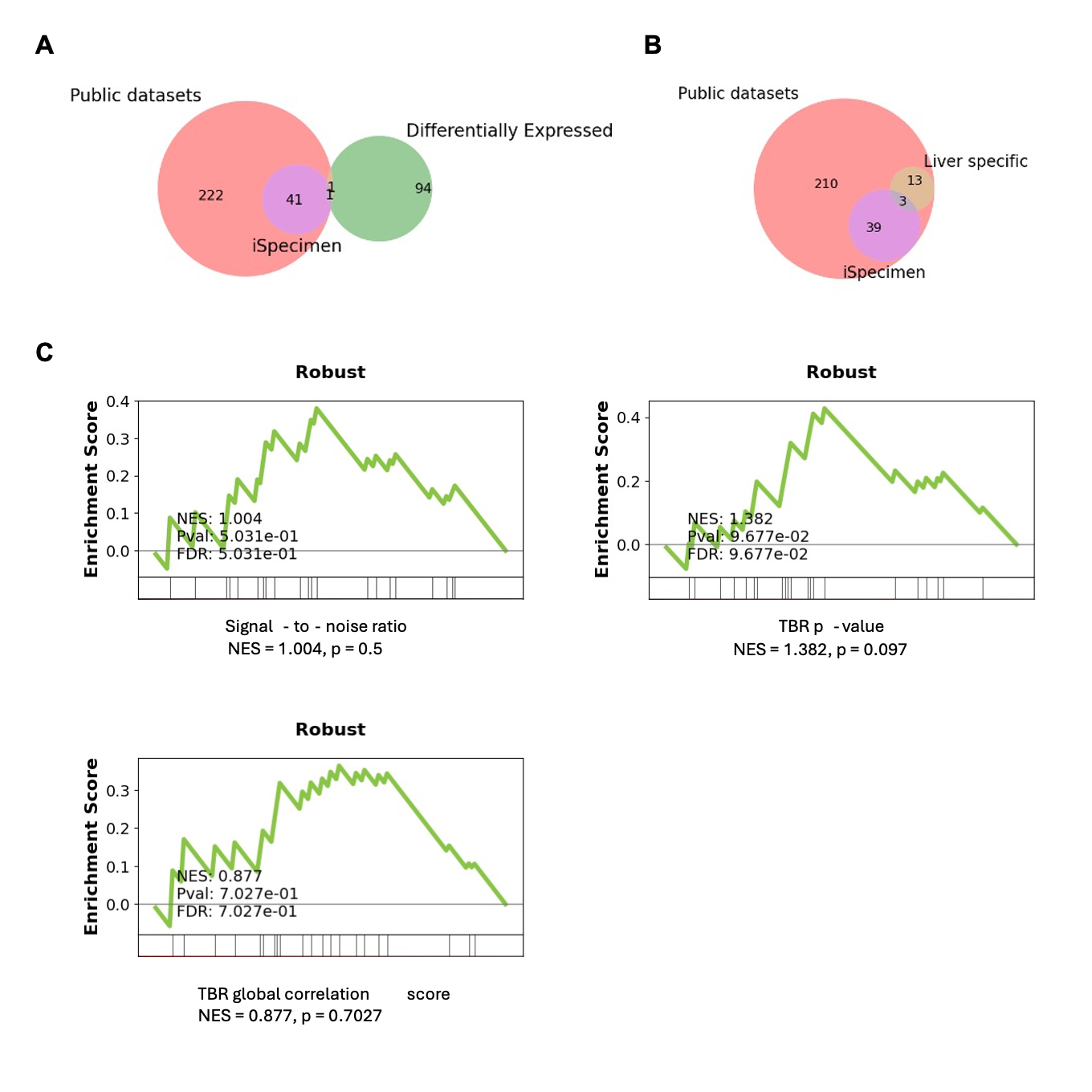

*Supplementary Figure 2:* ***(A) Differential gene expression, and (B) Cell-type specificity, are unable to identify generalizable blood-based biomarker candidates, (C) TBR signal-to-noise ratio, p-value, or correlation between gene expression profiles and the TBR disease trajectory do not enrich for generalizable biomarkers.***
